# Towards Generalizable Predictions for the Effects of Mutations on G-Protein Coupled Receptor Expression

**DOI:** 10.1101/2021.12.28.474371

**Authors:** Charles P. Kuntz, Hope Woods, Andrew G. McKee, Nathan B. Zelt, Jeffrey L. Mendenhall, Jens Meiler, Jonathan P. Schlebach

## Abstract

Missense mutations that compromise the plasma membrane expression (PME) of integral membrane proteins (MPs) are the root cause of numerous genetic diseases. Differentiation of this class of mutations from those that specifically modify the activity of the folded protein has proven useful for the development and targeting of precision therapeutics. Nevertheless, it remains challenging to predict the effects of mutations on the stability and/ or expression of MPs. In this work, we utilize deep mutational scanning data to train a series of artificial neural networks to predict the effects of mutations on the PME of the G-protein coupled receptor (GPCR) rhodopsin from structural and/ or evolutionary features. We show that our best performing network, which we term PMEpred, can differentiate pathogenic rhodopsin variants that induce misfolding from those that primarily compromise signaling. This network also generates statistically significant predictions for the effects of mutations on the PME of another GPCR (β_2_ adrenergic receptor) but not for an unrelated voltage-gated potassium channel (KCNQ1). Notably, our analyses of these networks suggest structural features alone are generally sufficient to recapitulate the observed mutagenic trends. Moreover, our findings imply that networks trained in this manner may be generalizable to proteins that share a common fold. Implications of our findings for the design of mechanistically specific genetic predictors are discussed.

## Introduction

Integral membrane proteins (MPs) are prone to misfolding and degradation, which leaves them vulnerable to the effects of destabilizing mutations.^1^ This intrinsic instability ultimately constrains their mutational tolerance.^2^ Indeed, a survey of 10,000 human genomes revealed that genetic variation is suppressed within regions encoding transmembrane domains (TMDs).^3^ Mutations that alter the physicochemical properties of TMDs can potentially disrupt membrane topology,^4^ folding,^5,6^ and/ or oligomerization.^7–9^ This type of mutation (class II) most often causes a reduction in the expression of the functionally mature protein at the plasma membrane (plasma membrane expression, PME), which is an underlying mechanism for numerous diseases of proteostasis.^1^ Efforts to target small molecules that correct such conformational defects to patients bearing class II variants have revolutionized the treatment of cystic fibrosis-^10–13^ a quintessential disease of MP misfolding.^14^ This breakthrough demonstrates that an understanding of the molecular effects of mutations can provide a decisive advantage in the development and targeting of precision therapeutics.^15^ Nevertheless, it remains a formidable challenge to rationalize the effects of most mutations on MP folding and function.^16^

According to the UniProt database, there are currently over 1,200 known disease-linked MPs, many of which are established drug targets.^17^ Some of these genes are associated with hundreds of genetic variations that elicit mechanistically diverse molecular effects that have been linked to one or more disease phenotypes. Most of these variants remain uncharacterized, and computational inference is therefore required to interpret their molecular effects. Most knowledge-based algorithms developed for such purposes (ie SIFT^18^ and PolyPhen2^19^) have been trained against collections of variants that have been loosely associated with a complex spectrum of clinical phenotypes (ie ClinVar). Though these predictors are generally useful, the performance is ultimately constrained by inconsistencies within the training data. Recent efforts generate more accurate predictions have instead turned to biochemical training data. A collection of 21,026 deep mutational scanning (DMS) measurements was recently used to develop a machine learning algorithm (Envision) that produces more accurate, quantitative predictions for the effects of missense variants on protein function.^20^ However, these training data were devoid of mutational data from MPs and the algorithm was not designed to distinguish between the effects of mutations on expression versus activity. The effect of mutations on the thermal stability of MPs^21^ or on their expression levels in *E. coli*^22,23^ have been predicted with some success, though it is unclear whether either of these factors should directly relate to PME levels in eukaryotic cells. Smaller sets of experimental measurements have been used to develop algorithms that predict the proteostatic, functional, or phenotypic effects within specific MPs of interest.^24–26^ Nevertheless, these specialized tools also exhibit limited accuracy and cannot produce generalized predictions for the effects of mutations within other proteins.

In the following, we describe the development of a computational tool to predict the impact of mutations on the PME of G-protein coupled receptors (GPCRs) that we term the PME predictor (PMEpred). Using a limited set of DMS measurements, we trained a series of artificial neural networks (ANNs) to predict the effects of mutations on the PME of the rhodopsin GPCR (Figure 1), the misfolding of which is associated with retinitis pigmentosa (RP) and a series of other related retinopathies.^27^ We show that PMEpred can differentiate class II rhodopsin variants from other disease variants that are known to primarily disrupt function. We also find that PMEpred generates statistically significant predictions for the impacts of mutations on the expression of the β_2_ adrenergic receptor (β_2_AR) GPCR, though not for an unrelated voltage-gated potassium channel (KCNQ1). These findings suggest that networks trained to recognize mutagenic trends in one receptor can be generalized to others that share the same fold. Finally, an analysis of the feature set used by these networks suggests mutagenic trends in PME can be recapitulated from structural features alone, and that evolutionary features offer relatively little predictive power. This finding highlights important considerations for the development of mechanistically specific genetic predictors.

**Figure 1.**
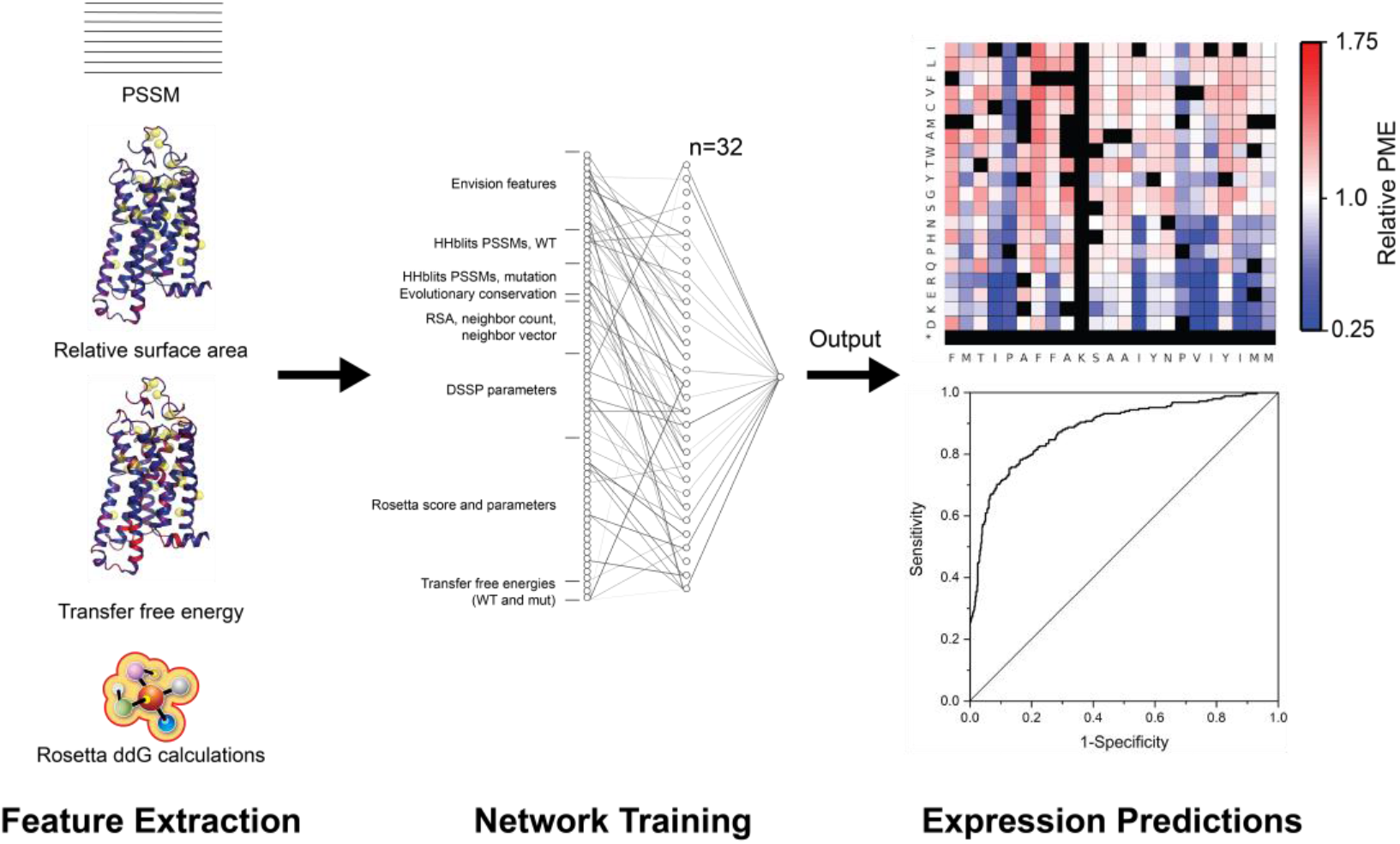
Training artificial neural networks to predict the plasma membrane expression of membrane protein variants. A schematic representation depicts the general training approach and network architecture used to develop the PMEPred algorithm.

## Results

### Feature Selection and Network Training

We recently utilized DMS to measure the relative PME of rhodopsin variants in HEK293T cells.^2^ In relation to other DMS results, these data are relatively limited in scope (808 mutations covering two TMDs). Nevertheless, to our knowledge, these data constitute the only example of a high-quality mutational scan that was specifically designed to measure the PME of MP variants within a human cell line. We therefore trained a series of ANNs consisting of 131 input nodes and a single hidden layer of 32 neurons to recapitulate the relative PME of these rhodopsin variants from an array of evolutionary and/ or structural features (Figure 1). Each ANN was trained using a previously described *k*-fold cross validation procedure in which the output of the network is iteratively refined against non-overlapping sub-sets of the training data.^24^ To avoid over-fitting, we utilized a dropout procedure in which neurons are occasionally ignored during the training process. We first trained an ANN using the feature set employed by Envision^20^ in addition to various other evolutionary and structural features (Table S1). To incorporate additional evolutionary information into the feature set, we derived evolutionary rates and a position-specific scoring matrix (PSSM) for rhodopsin. We also used a previously described structural model of human opsin to generate descriptors associated with the structural context of each mutated residue.^28^ To estimate the effects of each mutation on the free energy of the native conformation, we calculated Rosetta ddG values and incorporated each of the underlying energy terms as independent features. Finally, to account for any potential changes in the fidelity of cotranslational membrane integration,^28^ we used the Punta-Maritan scale to estimate the effects of mutations on the transfer free energy associated with the membrane integration of TMDs.^29^

An ANN trained with these features generates predicted relative PME values (PMEpred scores) that are correlated with the training data (Pearson’s R = 0.69, Figure 2A). Moreover, a receiver operating characteristic analysis suggests this network is effective in classifying variants based on whether they decrease the PME of rhodopsin (AUC = 0.873, Figure 2B). A sensitivity analysis reveals that the most predictive features include sequence-based descriptors (ie descriptors for residue properties) and characteristics related to secondary structure (ie DSSP terms, Table S2). It should be noted, however, that while evolutionary features are often the most powerful for functional prediction, we find that similar predictive power can be achieved using structural features alone (Figure S1, Table S3). Nevertheless, regardless of the specific features employed, we find this network architecture is generally capable of recapitulating the observed mutagenic trends. Hereafter, we will refer to the final version of this network that employs the full feature set as PMEpred.

**Figure 2.**
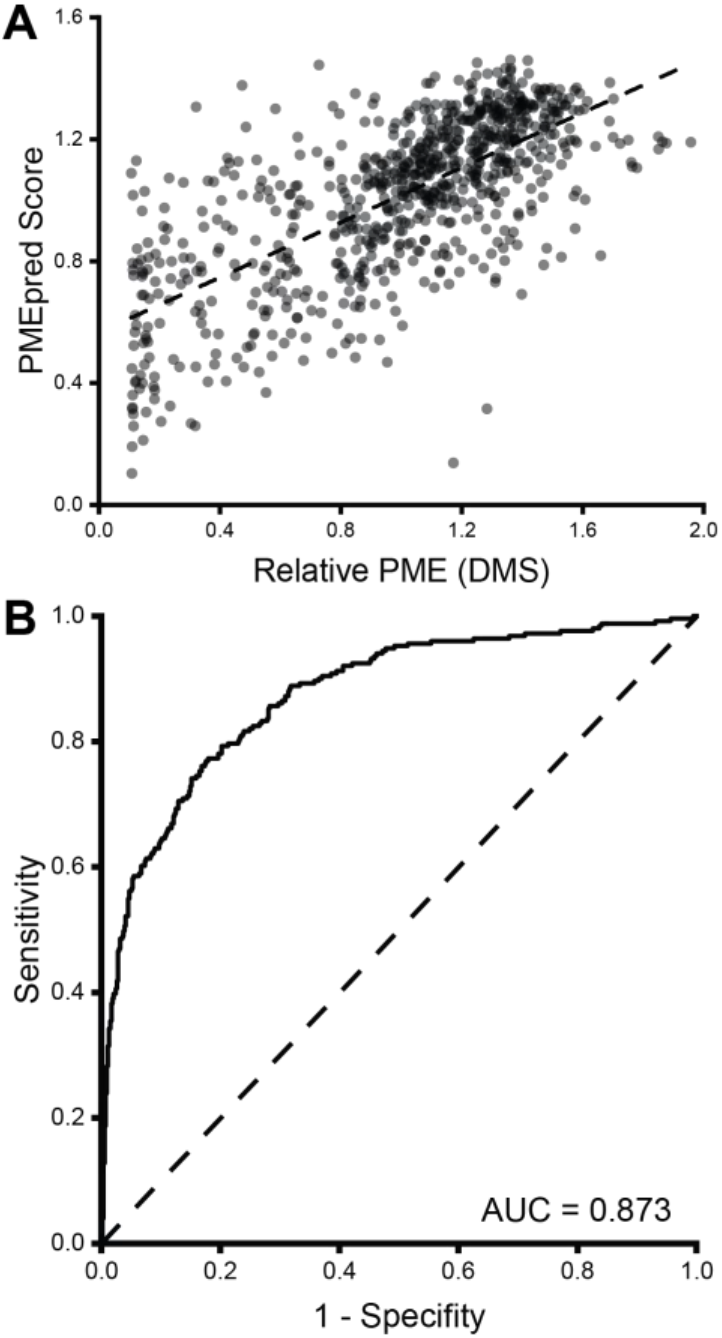
Evaluation of PMEpred scores in relation to experimental training data. A) The PMEpred scores for 808 missense variants of rhodopsin are plotted in relation to the corresponding deep mutational scanning data in the training set. A linear fit of the data is shown for reference (dashed line, Pearson’s R = 0.69). B) A receiver operating characteristic plot describes the ability of PMEpred scores to classify variants based on whether they decrease the PME of rhodopsin by at least 10% (Area Under the Curve = 0.873). The dashed line represents the position of a curve with zero predictive power.

### PMEpred Scores of Pathogenic Rhodopsin Variants

To assess whether PMEpred can differentiate misfolded variants from other loss-of-function (LOF) variants, we generated PMEpred scores for 128 mutations within rhodopsin that are known to cause RP or other related retinopathies. Though these disease variants are enriched with class II variants that compromise PME, the distribution of PMEpred scores for these variants is comparable to that of the comprehensive set of 6122 rhodopsin variants (Figure S2). We reason that this poor discrimination could stem from a lack of predictive power within the loop regions, which lack coverage in the training data. Indeed, when variants within the loops are omitted, leaving 2808 TMD variants, the PMEpred scores of RP variants skews lower than those of other TMD variants (Mann-Whitney p = 0.002, Figure S2). Moreover, the PMEpred scores for the known class II variants within this subset skew much lower than those that are known to specifically compromise function (Mann-Whitney p = 0.02, Figure 3). Overall, 71% of known class II variants are predicted to decrease PME (PMEpred score < 1.0, Matthew’s correlation coefficient = 0.39). The unclassified variants within this set exhibit a distribution of scores that is comparable to those of known class II variants (Figure 3), which is consistent with the fact that misfolded variants are generally the most common class of LOF variant.^5,6,30^ Together, these findings demonstrate that PMEpred can differentiate class II variants from other classes of mutations within TMDs with a 71% accuracy.

**Figure 3.**
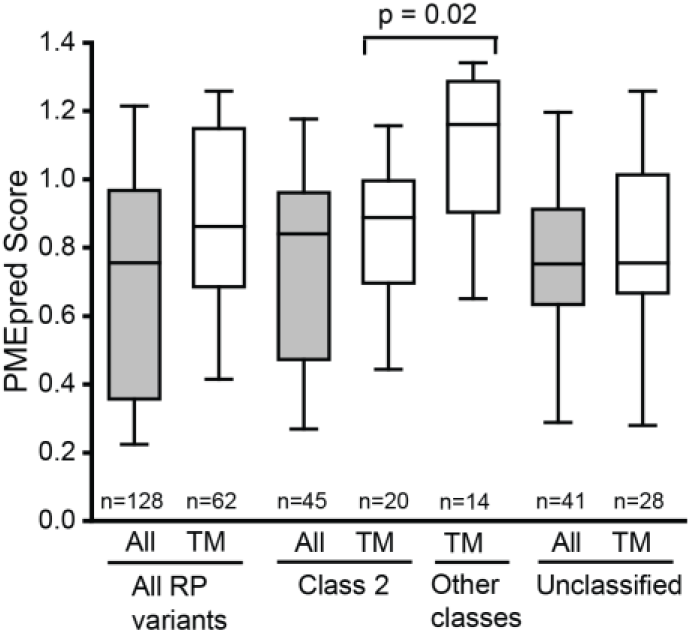
PMEpred Scores for known Rhodopsin Disease Variants. A box and whisker plot describes the distribution of PMEpred scores among all pathogenic missense variants within the TM domains of rhodopsin (second box) in relation to the subsets that are either known to induce misfolding (class II, fourth box), or are known to have normal expression (non-class II, fifth box), or are currently unclassified (seventh box). Gray boxes (first, third, and sixth) show scores for variants in loop regions in addition to TMDs. The center line reflects the median score. The upper and lower edges of the box reflect the positions of the 75^th^ and 25^th^ percentile. The upper and lower whiskers reflect the 90^th^ and 10^th^ percentile values. A p-value from a pairwise Mann-Whitney U-test comparing these distributions is shown.

### Evaluation of PMEpred Scores for Other Proteins

Given the limitations of the training data, we sought to determine whether PMEpred can generate useful predictions for other proteins. Benchmarking PMEpred is challenging given the current lack of other quantitative, large-scale data sets that specifically measure trends in PME. However, the functional effects of ~7,800 variants within the β_2_AR GPCR were recently reported.31 Though PMEpred is not designed for direct functional predictions, we would expect mutations that decrease PME to be enriched among LOF mutations. To evaluate PMEpred scores for LOF variants, we first compiled a β_2_AR feature set and used it to calculate PMEpred scores for each of its missense variants. We then applied a quality filter to remove noisy measurements from the experimental β_2_AR scan data and classified the remaining variants as either neutral or LOF based on the average functional score across all measured variants (Figure 4A). The distribution of PMEpred scores across the entire receptor are comparable for neutral and LOF variants (Figure 4B). However, LOF variants within TMDs exhibit significantly lower PMEpred scores relative to neutral TMD variants (Mann-Whitney p = 9.9 × 10^−14^, Figure 4B). This observation again suggests PMEpred is incapable of predicting the proteostatic effects of loop mutations, which we attribute to a lack of training data within these regions. Consistent with this interpretation, also observe discrimination between neutral and LOF variants within the smaller subset of variants within TMs 2 & 7 (Mann-Whitney p = 2.2 × 10^−5^, Figure 4B), which are the mutagenized helices included in the training data. Despite this limitation, our results show that PMEpred can differentiate between neutral and LOF of β_2_AR even though the later subclass is certain to contain variants that retain native expression levels. These observations demonstrate that networks trained with data from one GPCR are capable of spotting mutagenic trends in another.

**Figure 4.**
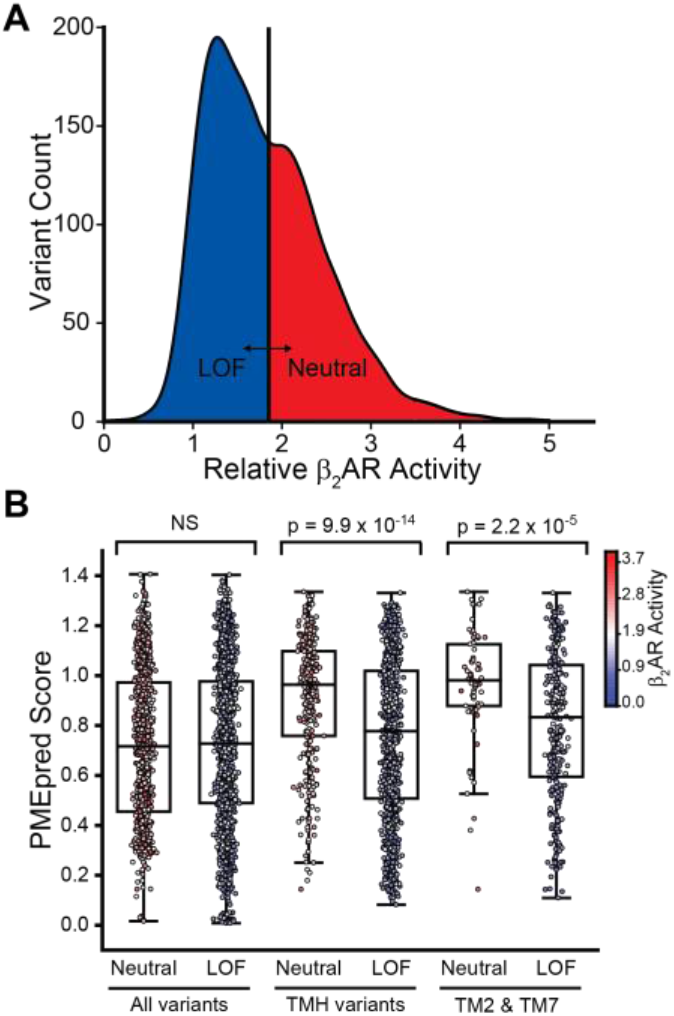
Distribution of PMEpred Scores for Neutral and Deleterious β_2_AR Variants. A) A histogram depicts the distribution of activity scores among 5,910 variants of the β_2_AR variants in the presence of 625 nM isoproterenol, as was reported in reference 31. A vertical line reflects 95% of the average functional score of all variants, which was used as a cutoff value to define the set of loss of function (LOF) variants. B) A box and whisker plot depicts the distribution of PMEpred scores among neutral and LOF variants within the entire set of variants (left), within the TMD residues (center), or within TM 2 & 7 (right). The center line in each box reflects the median score. The upper and lower edges of the box reflect the positions of the 75^th^ and 25^th^ percentile. The upper and lower whiskers reflect the 90^th^ and 10^th^ percentile values. p-values from Mann-Whitney U-tests comparing these distributions are shown for reference.

To evaluate the fidelity of PMEpred scores, we measured the PME of a set of 21 TM2 and TM7 variants that are predicted to increase or decrease the expression of β_2_AR (Table S4). Briefly, the β_2_AR variants bearing an N-terminal hemagglutinin (HA) epitope tag were transiently expressed in HEK293T cells. Surface immunostaining was then used to mark plasma membrane β_2_AR with a fluorescent anti-HA antibody. Cellular fluorescence intensities were then quantified by flow cytometry and normalized relative to that of cells expressing WT β_2_AR. Relative PME measurements do not appear to be linearly related to PMEpred scores (Figure 5A), which may arise in part from the difference in the dynamic range of PMEpred scores relative to normalized PME measurements. There also appear to be several false-negatives that score well yet are poorly expressed (Figure 5A). Despite these shortcomings, variants that are predicted to reduce expression by at least 10% (PMEpred score ≤ 0.9) exhibit significantly lower measured PME than those that are predicted to have higher expression (Mann-Whitney p = 0.01, Figure 5C) and the rank correlation between PMEpred scores and measured values is statistically significant (Spearman’s ρ = 0.49, p = 0.02). Trends are similar regardless of whether evolutionary features were included (Figure S3). These findings demonstrate that PMEpred can predict the proteostatic effects of mutations within the TMDs of a related GPCR.

**Figure 5.**
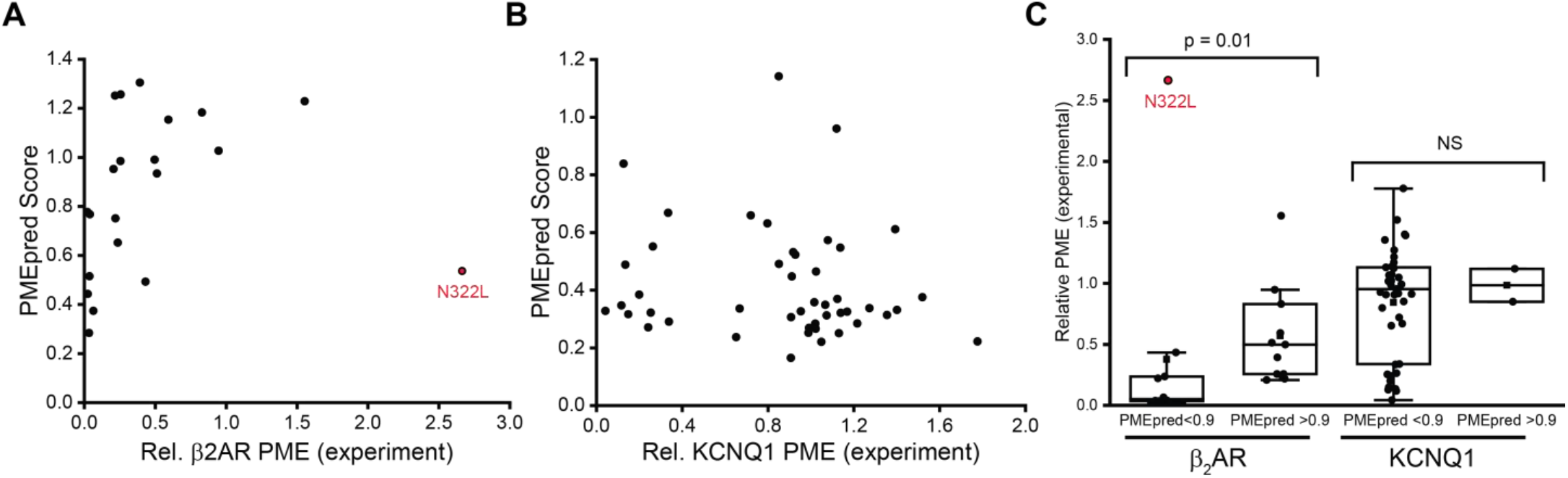
Evaluation of PMEpred scores in relation to Relative PME measurements of β_2_AR and KCNQ1 variants. A) The PMEpred scores for select β_2_AR variants are plotted against the corresponding measured value for the relative plasma membrane expression levels, which are normalized relative to that of WT. B) The PMEpred scores for select KCNQ1 variants are plotted against the corresponding measured value for the relative plasma membrane expression levels, which are normalized relative to that of WT. Measured values were previously reported in reference 6. C) A box and whisker plot compares the distributions of measured plasma membrane expression levels among β_2_AR (left) and KCNQ1 (right) variants that are predicted to decrease expression by at least 10% (PMEpred ≤ 0.9) to those for variants that are predicted to have robust expression (PMEpred>0.9). p-values from Mann-Whitney U-tests comparing these distributions are shown for reference.

We next sought to evaluate whether PMEpred can predict mutagenic trends within a structurally unrelated MP. For this purpose, we generated PMEpred scores for a set of 50 recently characterized pathogenic variants^6^ of the voltage gated potassium channel KCNQ1. Notably, nearly all of these variants are predicted to decrease the PME of KCNQ1 (Figure 5C), which is somewhat consistent with the fact that this set is enriched with class II variants.^6^ Nevertheless, PMEpred scores are generally uncorrelated with experimental PME measurements for these KCNQ1 variants (Figure 5B). These results imply PMEpred is unlikely to generate useful predictions for structurally divergent MPs.

## Discussion

Predicting the effects of mutations on the folding and/ or expression of MPs remains notoriously difficult,^16^ and efforts to improve such predictions address an imminent challenge in precision medicine.^15^ In this work, we used a limited set of DMS measurements to train an ANN to predict the PME of MP variants from various structural and evolutionary features. Though this is a relatively small set of training data (808 variants in two helices of a single receptor),^2^ we show that the resulting networks generate significant predictions for the proteostatic effects of known disease mutants within the other TMDs of the same protein (rhodopsin, Figure 3). We also find that this network can generate statistically significant predictions for the effects of TMD mutations on another related receptor (β_2_AR), but not for mutations within a structurally distinct ion channel (KCNQ1). The statistical power of our rhodopsin and β_2_AR predictions is generally comparable to that of current predictors of variant function that were developed with much larger sets of training data, which bodes well for the utility of DMS data for the development of more accurate expression predictors. We suspect the limitations exhibited by the current version of PMEpred, such as the inability to accurately predict effects within loop regions, can be overcome with the incorporation of more complete training data and features that more accurately capture the effects of mutations on protein stability. Nevertheless, it is unclear whether deeper training sets from a more diverse array of proteins will be sufficient to generate predictions that can be generalized across proteins with distinct folds.

The networks described herein were designed for a specific application-to predict the PME of MP variants in eukaryotic cells. Expression should be less challenging to predict than function or fitness, which are composites of expression and activity. Nevertheless, our efforts to predict these proteostatic effects revealed several observations that merit further consideration. First we note that, while evolutionary features are often potent for functional predictions,^32^ we find our networks exhibit comparable performance without them (Figure S1). This potentially suggests evolutionary data are biased towards function in a manner that ultimately undermines their utility for expression predictions. This is consistent with a recent analysis of DMS measurements on the effects of missense variants in PTEN and NUDT15, which shows that evolutionary rates are more closely associated with activity measurements whereas Rosetta ddG values correlate more closely with variant abundance.^33^ Our PME measurements of β_2_AR variants include one notable variant (N322L) that illustrates how evolutionary data may obscure the proteostatic effects of certain mutants (Figure 5 A & C). This mutation is heavily penalized due to the extreme conservation of N322, which forms part of the NPXXY motif. However, it also makes a relatively polar TMD more hydrophobic, which can dramatically enhance receptor PME.2,22,28 A second noteworthy observation concerns the low PME of several β_2_AR variants that are predicted to have robust expression (Figure 5A). Considering that most random mutations are destabilizing^34^ and that decreases in conformational stability often coincide with reduced PME,^5^ the general prevalence of high PMEpred scores among β_2_AR variants may hint at a more general issue (Figure 4B). The range of scores generated by these networks is dictated by those within the underlying training data from rhodopsin, which is known to be among the most stable GPCRs. While rhodopsin may be able to withstand the effects of most destabilizing mutations, the margin for error may be slimmer for a less stable protein like β_2_AR. The extent to which changes in stability (∆∆G) influence the folding efficiency and/ or PME of individual variants depends on the underlying free energy of folding (∆G) of the WT receptor- a critical unknown. Future algorithms may therefore benefit from approaches to account for this hidden variable. It may also be worth considering whether this factor may contribute to protein-specific variations in performance among leading functional predictors.^20^

The investigations described herein provide new tools for the prediction of the PME of GPCR variants while highlighting important considerations for the development of mechanistically specific prediction algorithms. Though the predictions generated by the current version of PMEpred are not necessarily accurate enough to call individual variants, they could potentially be used to help guide protein engineering efforts that leverage massively parallel gene synthesis. We expect the performance of such predictions to continue to improve as additional training data become available. Performance may also be improved by approaches to improve the curation of biased feature sets.

## Materials & Methods

### Curation of Feature Sets

An initial set of features was derived from those previously employed by Envision (listed in Supplementary Table 2 of Reference 20),^20^ except for those that cannot be derived obtained from comparative structural models (i.e. B-factor). We added several additional features to the final set. Features included the relative solvent accessible surface areas for each residue, which was calculated from our structural models using NACCESS. We included other features related to solvent accessibility such as neighbor count and neighbor vector, which we calculated using the Biochemical Library (BCL).^35^ Finally, the depth of an amino acid as measured by the distance to the surface of the protein was obtained using the Depth web server.^36^ Features related to secondary structure and hydrogen bonding were obtained from the program DSSP.^37^ We used HHblits^38^ to generate a series of PSSMs with varying stringencies to serve as independent features that describe how individual substitutions relate to the observed mutational tolerance. PSSMs were derived from rhodopsin sequences mined from the Uniprot20 database at e-value thresholds 1e^−2^, 1e^−3^, 1e^−4^, 1e^−5^, and 1e^−10^. Our feature set also included evolutionary rates for each position that were derived with Consurf using the uniref90 database with PSI-BLAST and an e-value threshold of 0.0001 (four iterations).^39^ Sequences that were more than 95% similar to one another were eliminated using CD-HIT.^40^ 468 sequences in total were identified and aligned using MAFFT.^41^ To incorporate data from human genomes/ transcriptomes, the missense tolerance ratio for each residue in human rhodopsin was included as an independent feature.^42,43^ To integrate information related to conformational dynamics, we used the DynaMut server to carry out normal mode analysis on our model of human opsin and generated features for the calculated mean fluctuation and deformation at each position.^44^ Transfer free energies for wild-type and mutant amino acids were derived from the knowledge-based scale of Punta and Maritan.^29^ We also included previously reported descriptors related to the physicochemical properties of amino acid side chains such as volume, polarity, and hydrophilicity.^45^ Unweighted energy terms and composite scores derived from Rosetta ddG calculations were included as independent features. Feature sensitivity analysis was performed by calculating the L1 norm of the signal associated with each feature.^46^ Each row within the dataset represents single variants rather than single positions within the sequence.

### Network Training

ANNs were implemented in bcl: model: Train within the Biochemical Library (BCL), a code repository maintained primarily by the Meiler laboratory at Vanderbilt University (http://www.meilerlab.org/index.php/bclcommons/show/b_apps_id/1) . The neural network had a single hidden layer with 32 neurons. The transfer function employed was the sigmoid function. The momentum employed was 0.3 and the learning rate was 0.05. Dropout was used, and the probability of dropout in the input layer was 25%, while the probability of dropout in the hidden layer was 50%. Each network was trained for 10,000 iterations.

### Plasmid Preparation and Mutagenesis

PME measurements were carried out using a transient expression vector we created by inserting a synthetic gene block encoding the cDNA sequence of the human β_2_AR (Integrated DNA Technologies, Coralville, IA) into a modified pcDNA5 FRT using the NEBuilder kit (New England Biolabs, Ipswitch, MA). The final construct was designed to generate β_2_AR bearing an N-terminal influenza hemagglutinin tag from a transcript that also features a downstream internal ribosome entry site (IRES)-dasher GFP cassette that facilitates the unambiguous identification of positively-transfected cells. Mutations of interest were introduced into the β_2_AR cassette by site-directed mutagenesis. Plasmids used for transient expression in HEK293T cells were prepared using the Zymopure Midiprep Kit (Zymo Research, Irvine, CA).

### Plasma Membrane Expression Measurements

The relative PME of β_2_AR variants was measured by flow cytometry using a simplified version of a previously reported method.5,28 Briefly, β_2_AR variants were transiently expressed in HEK293T cells using Lipofectamine 3000 (Invitrogen, Carlsbad, CA). Transfected cells were then harvested 2 days post transfection using TrypLE (Gibco, Grand Island, NY). Intact cells were then immunostained with a DyLight 550-conjugated anti-HA antibody (Invitrogen, Carlsbad, CA) for 30 minutes in the dark at room temperature. Following surface immunostaining, the cells were fixed using solution A from the Fix and Perm kit (Invitrogen, Carlsbad, CA) then washed twice and resuspended in phosphate buffered saline containing 2% fetal bovine serum. Cells were then filtered to remove aggregates prior to the analysis of cellular fluorescence profiles using a BD LSRII flow cytometer (BD Biosciences, Franklin Lakes, NJ). Cellular fluorescence profiles were analyzed using FlowJo (Treestar, Ashland, OR). The mean DyLight 550 intensity of several thousand positively transfected cells expressing β_2_AR variants, which were identified based on bicistronic dasher GFP expression, was normalized by that of cells expressing WT β_2_AR to derive relative PME values. Reported values reflect the average value from three biological replicates.

## Acknowledgments

We thank Hui Huang, Georg Kuenze, and Charles R. Sanders for providing expression data and structural models for KCNQ1. We thank Christiane Hassel and the Indiana University Flow Cytometry Core Facility for technical support. This research was supported in part by a grant from the National Institutes of Health (NIH) (R01GM129261 to J.P.S.).

