## Supplemental Material for "Towards Generalizable Predictions for the Effects of Mutations on G-Protein Coupled Receptor Expression"

*This File Includes:*

Figure S1
Figure S2
Figure S3
Table S1
Table S2
Table S3
Table S4

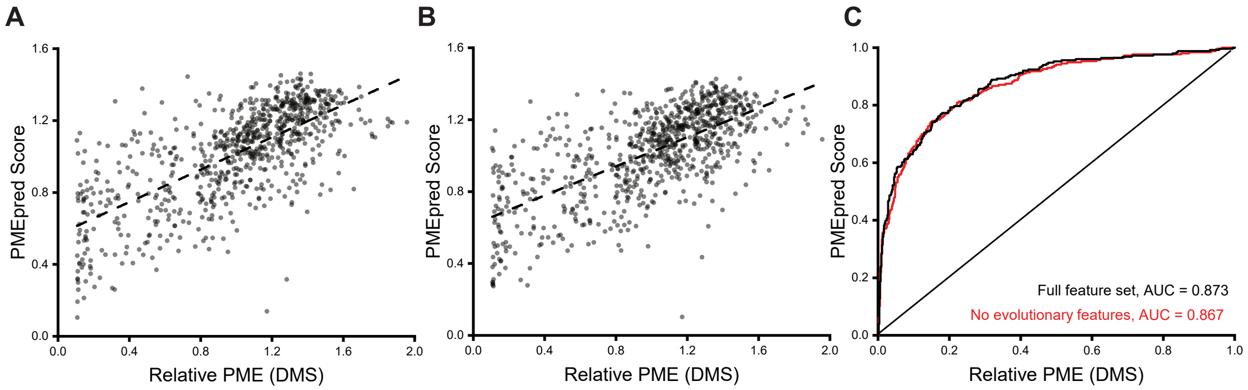


**Figure S1. Effect of evolutionary features on the performance of PMEPred.** The performance of neural networks trained with distinct feature sets is compared. A) PMEpred scores derived from a network trained with the full feature set are plotted against corresponding DMS measurements. A linear fit is included for reference (Pearsons R= 0.69) B) PMEpred scores derived from a network trained with a feature set that exludes any relating to sequence evolution are plotted against corresponding DMS measurements. A linear fit is included for reference (R= 0.67). C) Receiver-operating characteristic (ROC) curves for versions of the neural network trained with the full feature set (black) or with non-evolutionary features (red) are shown along with the corresponding values for the area under each curve (AUC).

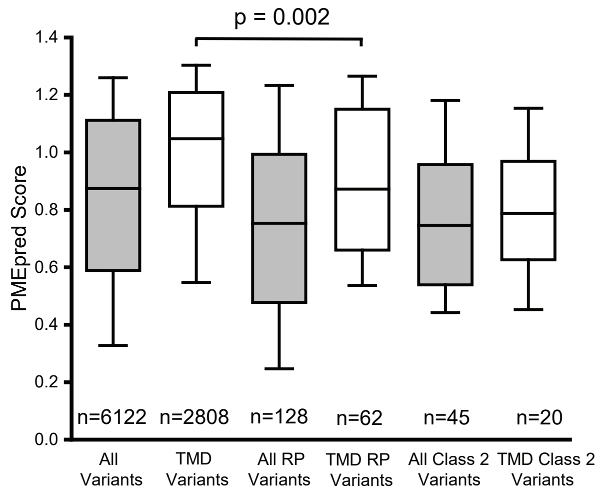


**Figure S2. Comparison of PMEpred scores for retinopathy variants to all possible rhodopsin variants.** PMEpred scores were extended to the full set of rhodopsin missense variants by excluding Rosetta ddG values, which could not obtained for all variants, from the feature set. The distribution of PMEpred scores for among all rhodopsin variants (first box) are compared to various subsets of variants including those within TM domains (second box), all known retinopathy variants (third box), retinopathy variants within TM domains (fourth box), all class II retinopathy variants (fifth box), and retinopathy variants within TM domains (sixth box).

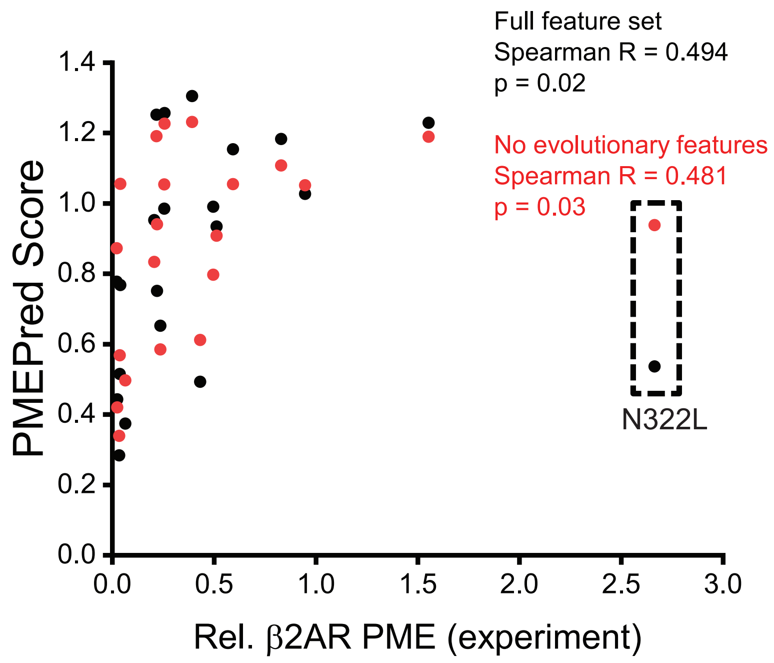


**Figure S3. Effect of evolutionary features on PMEPred Predictions for $\beta_2$AR .** PMEpred predictions for β2-adrenergic receptor variants generated by versions of the network trained with the full feature set (black) or without evolutionary features (red) are plotted against corresponding variant PME measurements. Spearman rank correlation coefficients and the corresponding p-values for each set are shown for reference. The points corresponding to the N322L outlier are indicated within the dashed box for reference.

**Supplementary Table 1. Annotated List of Machine Learning Features.**

| Feature name | Description |
|---|---|
| hhblits_1e-2_var, hhblits_1e-2_wt, hhblits_1e-3_var, hhblits_1e-3_wt, hhblits_1e-4_var, hhblits_1e-4_wt, hhblits_1e-5_var, hhblits_1e-5_wt, hhblits_1e-10_var, hhblits_1e-10_wt | HHBlits profile scores for wild-type and mutant amino acids<br>e-value thresholds: $10^{-2}$, $10^{-3}$, $10^{-4}$, $10^{-5}$, $10^{-10}$ |
| consurf_score | Rate of evolution derived from Consurf |
| rsa_whole_res, rsa_sidechain, rsa_mainchain, rsa_nonpolar, rsa_polar | Relative surface area from NACCESS |
| nco_0, nco_3 | Neighbor count (at least 0 or 3 residues away in primary sequence) |
| nvector | Neighbor vector |
| dssp_NHO1_part, dssp_NHO1_elec, dssp_OHN1_part, dssp_OHN1_elec, dssp_NHO2_part, dssp_NHO2_elec, dssp_OHN2_part, dssp_OHN2_elec, dssp_TCO, dssp_kappa, dssp_alpha, dssp_phi, dssp_psi | Secondary structure features from DSSP |
| rosetta_score | Rosetta ddG score for mutation |
| dslf_ca_dih, dslf_cs_ang, dslf_ss_dih, dslf_ss_dst, fa_atr, fa_dun, fa_intra_rep, fa_mpenv, fa_mpenv_smooth, fa_mpsolv, fa_pair, fa_rep, hbond_bb_sc, hbond_lr_bb, hbond_sc, hbond_sr_bb, omega, p_aa_pp, pro_close, rama, ref | Unweighted terms from Rosetta ddG calculations |
| depth_allatom, depth_mainchain | Residue depth within structure |
| mtr | Missense tolerance ratio (MTR) for position |
| nma_deform, nma_fluct | Fluctuation and deformation calculated from DynaMut |
| wt_maritan_dg, mut_maritan_dg, del_maritan_dg | Punta-Maritan transfer free energies, mutant and wild-type |
| wt_charge, mut_charge, del_charge | Residue charge at pH 7.4, wild-type and mutant |
| wt_pol, mut_pol, del_pol, wt_vol,mut_vol, del_vol, wt_hyd, mut_hyd, del_hyd | Residue volume, hydrophilicity, and polarity |
| AA1_pI, AA2_pI, deltaPI, AA1_weight, AA2_weight, deltaWeight, AA1_volume, AA2_volume, deltavolume, Grantham, AA1_PSIC, AA2_PSIC, delta_PSIC, mut_msa_congruency, seq_ind_closest_mut, evolutionary_coupling_avg, evolutionary_coupling_avg_norm | Features from ENVISION |

**Supplementary Table 2.** Feature scores derived from the L1 norm of signal associated with each input. (network trained with all features)

| Feature | Feature score |
|---|---|
| dssp_NHO1_part | 2.09551 |
| wt_charge | 0.945401 |
| dssp_NHO2_part | 0.929787 |
| dssp_NHO1_elec | 0.806575 |
| mut_maritan_dg | 0.751033 |
| fa_pair | 0.583702 |
| seq_ind_closest_mut | 0.488305 |
| wt_vol | -0.47271 |
| AA1_volume | -0.4592 |
| wt_hyd | -0.45183 |
| hhblits_1e-4_wt | -0.44561 |
| dssp_psi | 0.441506 |
| dslf_ss_dst | -0.42359 |
| nvector | -0.41933 |
| pro_close | -0.41892 |
| nco_0 | 0.404513 |
| AA1_pI | -0.39424 |
| wt_pol | -0.39204 |
| dslf_cs_ang | 0.389233 |
| dssp_OHN2_elec | -0.3867 |
| fa_mpenv_smooth | 0.379692 |
| hhblits_1e-5_wt | -0.37906 |
| deltaPI | -0.37296 |
| hhblits_1e-10_wt | -0.3725 |
| del_maritan_dg | 0.371219 |
| fa_dun | 0.36911 |
| AA2_PSIC | -0.36288 |
| mut_msa_congruency | -0.35236 |
| hhblits_1e-2_wt | 0.346712 |
| hbond_bb_sc | -0.33974 |
| del_pol | -0.32635 |
| hbond_lr_bb | -0.32363 |
| del_hyd | -0.32014 |
| rama | -0.31727 |
| hhblits_1e-3_wt | 0.314568 |
| fa_intra_rep | 0.314018 |
| nma_deform | -0.30294 |
| rsa_mainchain | 0.301172 |
| depth_mainchain | 0.300738 |

| | |
|---|---|
| p_aa_pp | -0.29638 |
| AA1_weight | -0.289 |
| dslf_ss_dih | -0.27788 |
| AA2_weight | 0.275559 |
| nco_3 | 0.270647 |
| fa_atr | 0.269184 |
| hbond_sc | -0.25667 |
| fa_rep | -0.25534 |
| wt_maritan_dg | -0.24949 |
| rsa_polar | -0.23814 |
| mut_vol | 0.232909 |
| del_charge | -0.22654 |
| delta_PSIC | -0.21742 |
| mtr | -0.21588 |
| dssp_TCO | 0.214118 |
| deltaWeight | 0.21179 |
| omega | -0.21126 |
| Grantham | 0.206348 |
| dssp_OHN1_part | -0.20586 |
| mut_charge | -0.19917 |
| depth_allatom | 0.198595 |
| deltavolume | -0.19359 |
| mut_hyd | -0.19095 |
| dssp_kappa | -0.18797 |
| dssp_OHN1_elec | 0.185889 |
| hhblits_1e-2_var | -0.18577 |
| nma_fluct | -0.18172 |
| mut_pol | -0.17955 |
| hhblits_1e-3_var | -0.17566 |
| ref | -0.1738 |
| evolutionary_coupling_avg | 0.165515 |
| evolutionary_coupling_avg_norm | 0.162518 |
| fa_mpsolv | -0.16036 |
| rosetta_score | 0.158463 |
| fa_mpenv | -0.15523 |
| dslf_ca_dih | 0.153994 |
| consurf_score | -0.15134 |
| dssp_NHO2_elec | -0.13638 |
| dssp_phi | 0.125933 |
| AA1_PSIC | -0.11592 |
| hhblits_1e-5_var | 0.099863 |

| | |
|---|---|
| hbond_sr_bb | -0.09692 |
| dssp_alpha | 0.093768 |
| hhblits_1e-10_var | 0.085289 |
| AA2_pI | -0.07802 |
| dssp_OHN2_part | -0.07343 |
| rsa_nonpolar | -0.07045 |
| rsa_whole_res | -0.05939 |
| hhblits_1e-4_var | 0.046248 |
| AA2_volume | 0.024585 |
| del_vol | 0.021995 |
| rsa_sidechain | 0.003441 |

**Supplementary Table 3.** Feature scores derived from the L1 norm of signal associated with each input. (network trained without evolutionary features)

| Feature | Feature score |
|---|---|
| wt_vol | -0.48648 |
| AA1_volume | -0.48237 |
| pro_close | -0.46114 |
| nma_deform | -0.42335 |
| dssp_OHN2_elec | -0.40463 |
| AA1_pI | -0.39753 |
| wt_hyd | -0.38519 |
| dslf_ss_dst | -0.37719 |
| hbond_lr_bb | -0.37092 |
| AA1_weight | -0.3471 |
| mtr | -0.33919 |
| dssp_NHO2_elec | -0.33819 |
| p_aa_pp | -0.30276 |
| hbond_sc | -0.29516 |
| hbond_sr_bb | -0.29435 |
| rama | -0.29359 |
| wt_pol | -0.28734 |
| mut_charge | -0.28497 |
| rsa_polar | -0.2771 |
| del_pol | -0.27487 |
| del_hyd | -0.26556 |
| fa_rep | -0.26497 |
| hbond_bb_sc | -0.25774 |
| deltaPI | -0.25644 |
| dssp_OHN1_part | -0.2419 |
| deltavolume | -0.23667 |
| ref | -0.23162 |
| dssp_kappa | -0.21993 |
| omega | -0.20462 |
| nvector | -0.19677 |
| dssp_phi | -0.19589 |
| fa_mpsolv | -0.17259 |
| rsa_nonpolar | -0.16676 |
| wt_maritan_dg | -0.15867 |
| AA2_pI | -0.15735 |
| rsa_whole_res | -0.12919 |
| fa_mpenv | -0.09121 |
| dslf_ss_dih | -0.09091 |
| rsa_sidechain | -0.07336 |

| | |
|---|---|
| del_vol | -0.07274 |
| dssp_OHN1_elec | -0.07224 |
| nma_fluct | -0.04615 |
| dssp_OHN2_part | -0.04503 |
| mut_hyd | -0.026 |
| del_charge | -0.02349 |
| AA2_volume | -0.01241 |
| mut_pol | -0.0071 |
| dslf_ca_dih | 0.061521 |
| dssp_alpha | 0.095689 |
| deltaWeight | 0.110748 |
| depth_allatom | 0.133895 |
| fa_intra_rep | 0.150044 |
| mut_vol | 0.150974 |
| rsa_mainchain | 0.203723 |
| AA2_weight | 0.205541 |
| depth_mainchain | 0.227691 |
| nco_3 | 0.231446 |
| fa_atr | 0.247793 |
| rosetta_score | 0.26451 |
| del_maritan_dg | 0.294545 |
| Grantham | 0.305201 |
| nco_0 | 0.359271 |
| dssp_TCO | 0.377142 |
| dslf_cs_ang | 0.379034 |
| fa_mpenv_smooth | 0.389759 |
| dssp_psi | 0.411037 |
| dssp_NHO1_elec | 0.414084 |
| fa_dun | 0.562894 |
| fa_pair | 0.669426 |
| mut_maritan_dg | 0.772402 |
| dssp_NHO2_part | 0.944911 |
| wt_charge | 1.24466 |
| dssp_NHO1_part | 1.83275 |

**Supplementary Table 4.** List of ADRB2 variants tested

| Variant | Activity | PME (exp) | Score (Full set) | Score (Non-evol. set) |
|---|---|---|---|---|
| I72K | 1.28 | 0.22 | 0.75 | 0.94 |
| S74F | 1.03 | 0.04 | 0.77 | 1.06 |
| C77M | 2.07 | 0.95 | 1.03 | 1.05 |
| A78L | 1.19 | 0.26 | 1.26 | 1.23 |
| V81D | 1.02 | 0.06 | 0.37 | 0.50 |
| V87S | 1.18 | 0.22 | 1.25 | 1.19 |
| V87Q | 1.26 | 0.21 | 0.95 | 0.83 |
| F89W | 1.98 | 0.51 | 0.93 | 0.91 |
| F89H | 1.41 | 0.43 | 0.49 | 0.61 |
| A91T | 2.05 | 0.50 | 0.99 | 0.80 |
| A91R | 0.93 | 0.03 | 0.28 | 0.34 |
| L310V | 2.74 | 0.59 | 1.15 | 1.05 |
| L311Q | 1.53 | 0.02 | 0.78 | 0.87 |
| N312S | 0.92 | 0.83 | 1.18 | 1.11 |
| W313K | 1.16 | 0.04 | 0.52 | 0.57 |
| I314D | 1.09 | 0.03 | 0.44 | 0.42 |
| F321I | 1.89 | 0.26 | 0.99 | 1.05 |
| F321Y | 1.65 | 0.24 | 0.65 | 0.58 |
| N322L | 1.32 | 2.67 | 0.54 | 0.94 |
| Y326C | 1.12 | 1.55 | 1.23 | 1.19 |
| Y326N | 1.90 | 0.39 | 1.30 | 1.23 |
